# Temporal dynamics and policy implications in an innovation arms race

**DOI:** 10.1101/2025.02.13.638049

**Authors:** Michael Chimento, Edwin Dalmaijer, Barbara C. Klump, Lucy M. Aplin

## Abstract

Human-modified environments offer novel resources, but their exploitation can be a source of human-wildlife conflict. Residents of Sydney have reported increasing cases of bin-opening behavior by sulphur-crested cockatoos (*Cacatua galerita*), with evidence this behavior is socially learned between birds. Households protected their bins, yet cockatoos have learned to defeat these defences. In response, residents increased their defence-level, setting the stage for a behavioral “arms race”. Here, we investigate this arms race by combining field observations with agent-based modeling. We systematically document protections in a suburban locality over two years, revealing spatial clustering of protections indicative of social learning among residents. We find that protections have decreased since 2019, and overly-costly protection is dis-preferred. With a controlled assay, we characterize the knowledge of local cockatoos, finding differential proficiency in defeating protections. Surprisingly, we identified several cockatoos that can defeat high-efficacy protections, such as locks. Finally, we simulate interactions between two populations of learning agents representing households and cockatoos. We find that social learning accelerates adaptation in both species, while coordination reduces costs associated with defensive escalation. However, policy interventions can have unintended consequences, accelerating cockatoos’ skill acquisition and shifting conflict to neighboring areas. Our findings highlight the dynamic nature of human-wildlife conflict and the importance of considering behavioral feedback loops in urban wildlife management.

## 1 Introduction

Innovation and cultural adaptation have uniquely positioned humans as a dominant ecological force on our planet (1). Ever growing human land-use change can disturb and displace local wildlife, but may also provide attractive, novel resources which can benefit some other animal species (2; 3). This is most dramatically observed in cities, where urban environments are highly novel and can provide an entirely new suite of resources of those species able to colonise them (2; 4). In addition to this highly novel environment, urban areas have a high density of humans. This leads to the close co-occurrence of humans and other urban-adapted animals, and the potential for both positive and negative human-wildlife interactions (5). Although such interactions have always been a defining feature of human existence, contemporary research interest in human-wildlife conflict has exponentially increased over the past two decades (6), especially given its economic and biological costs, and its relevance in the Anthropocene.

Human-wildlife conflict encompasses many types of interactions, but can broadly be defined as actions by humans or wildlife that have an adverse effect on the other, and the perception that wildlife threatens human safety, health, or property (7; 8; 6). Traditionally these conflicts have mostly occurred in primary production contexts, with well-known examples including the destruction of crops by African and Asian elephants (*Loxodonta africana* and *Elephans maximus*) (9), the predation of livestock by grey wolves (*Canis lupus*) (10), the depredation of fish from lines and nets by cetaceans (11), and raiding of bee-hives and orchards by brown bears (*Ursus arctos*) (12). But examples in urban contexts are increasingly reported, including house break-ins by chacma baboons (*Papio ursinus*) (13), klepto-parasitism of humans by gulls (*Larus spp.*) (14), and depredation of pets by coyotes (*Canis latrans*) (15). However, conflict need not be dramatic or violent, and there are innumerable nuisance conflicts between humans and animal, especially in urban contexts and involving human food waste (16; 17; 5).

Accessing food waste may be particularly cognitively challenging if the waste is not in the open, requiring some combination of neophilia, individual innovation and social learning, and behavioural flexibility (18; 19). In cases where food-waste is protected, it can become an extractive foraging problem, requiring behavioral innovations to overcome protections, and memory to keep those innovations in repertoire. In particular, social learning can provide a shortcut to adaptive information (20; 21; 22), and likely plays a large role in the spread of nuisance behaviors through animal populations (23). However, for social learning to take place, a behavior must first be innovated through individual learning. Particularly habituated and neophilic individuals may interact more with anthropogenic objects and structures, leading to behavioral innovations that can be used to exploit novel resources (24; 25; 26), such as the case of sugar-packet opening by noisy miners (*Manorina melanocephala*) or milk-bottle cap opening by great tits (*Parus major*) (27; 28). Even if only a small percentage of a population is capable of innovation, social learning can greatly increase the number of individuals using such innovations.

One contemporary example of a nuisance conflict is the bin-opening behavior of urban-living sulphur crested cockatoos (*Cacatua galerita*, SCC henceforth) in New South Wales, Australia. SCC are widespread across eastern Australia and have long been known as agricultural pests, feeding on cereal crops and orchard fruits (29; 30). They have also adapted to urban habitats where they adopt a generalist diet (31). In southern Sydney this generalist diet now includes food from household rubbish bins, whereby individual lift and flip bin-lids to scavage various food waste items such as bread. Klump et al. (32) identified the geographic spread of bin-opening behaviour from three suburbs to over forty suburbs over the course of several years. A network-based diffusion analysis of its spread (33; 34), along with spatial autocorrelation of opening techniques provided evidence for the social transmission of bin-opening behavior, meaning that SCC are likely learning how to exploit this resource from each other (35).

Litter left from SCC bin-opening behaviour has become a general nuisance for residents of communities where it is occurring, leading residents to implement a variety of protection measures, such as heavy bricks, ropes, plastic and purpose-designed locks (36). Several local councils have even implemented free bin-lock programs available to residents who are experiencing conflict with SCC. A follow-up study created a taxonomy of bin protections (36). Spatial autocorrelation in protection measures was observed, suggesting that residents were influenced by their neighbors’ protection choices. Field observations of opening behavior found that cockatoos could defeat low-efficacy options, such as loose rocks, and found no cases of birds defeating higher-efficacy protections, such as locks. Altogether, this led the authors to suggest that an “innovation arms race” was potentially occurring between cockatoos and residents, whereby both species were learning and modifying their behaviour in response to the behaviour of the other (36).

In order to explore this arms race variant of human-wildlife conflict in more detail and from a longitudal perspective, we recorded bin protection behaviour by households in two consecutive winters in one of three “hotspot” suburbs from which bin-opening behaviour is believed to have originated (32). Using transects, we recorded direct observations of bin-opening each week and matched this with an experiment testing the knowledge state of individually-marked SCC to better understand the distribution of birds’ knowledge about how to open bins and defeat protections. Finally, we used this empirical data to inform the design of an agent based model that simulates human-wildlife conflict between competing populations of learning agents. We use this model of a behavioral arms races to explore how costs and benefits to residents change in response to whether behaviors are individually or socially learned (either by residents or SCC) and whether localities coordinate their strategies. In doing so, the model also highlight critical factors for simulation outcomes that may be of future interest for research, such as the memory depth and movement decision-making strategies of SCC.

## 2 Methods

### 2.1 Transect observations of bin protections

During the winter months of 2023 (July 31 to October 27, 2023) and 2024 (July 1 to September 6, 2024) we collected information on bin protections used by residents of a coastal locality in north Wollongong that is surrounded by national park and state forest. In this area, people generally wheel their bins to the street-curb the evening before a weekly bin-collection day, making them available for council collection trucks in the morning, but also making them vulnerable to cockatoos. We conducted transects that took place on the evening prior to, and morning of, household bin collection, and recorded the location of all observed bins. Based on the existing taxonomy (36), we identified protection methods and their associated efficacies: block (placement of bin such that full opening is blocked), loose and fixed weight, rim (plastic around rim to increase difficulty of balancing), spikes, stick, bottle (through the hinges), stick (through the hinges), spring, rope, wire, bungee, hook, and lock. Bins with more than one defence (e.g. bottle and weight) were coded as multiply defended. Bins with multiple of the same defence (e.g. 2 weights) were coded as multiply defended. For bins with multiple protections, the non-broken defence of highest efficacy was assigned. A complete list of all combinations of defences can be found in Table S2. Example photos of common defence types can be found in Figure S1.

We predicted that if local residents were socially learning from neighbours, bin protections should show evidence of geographic clustering. To test this, we computed geodesic distance between bins using the sf pack-age (37), and computed driving distance between bins using the openrouteservice package (38). Using both inverse geodesic distance and inverse driving distance networks, we measured assortativity using the assortnet package (39) of: 1) whether or not a bin was protected (binary measure), 2) protection efficacy (continuous measure), and 3) protection method (ordered factor). P-values for assortment were computed using 1000 permutation tests. Less than 9% of households used less common protection methods (strategic placement of bins, plastic rims, bungees, wire, hooks, ropes, and springs), and these were pooled into an “other” category for analysis. Protection methods for bins with multiple protections were pooled as a single “multi-protection” category.

### 2.2 Observations of bin opening behaviour by SCC

#### 2.2.1 Study population

In 2024, we temporarily habituated and marked 193 SCC nearby to the focal SCC night roost with uniquely identifiable color combinations, following established protocol (32; 40). During marking, we collected a feather sample where possible, and use morphometrics to assign 81 individuals’ age and sex of adults based on eye color. For 89 individuals, we were able to determine sex from DNA extracted from the feather sample. Combining these data sources, we were left with a dataset with 121 marked individuals with known age and/or sex, summarized in Table 1. All marking, feather collection and experiments were conducted under approval by the Australian National University Animal Ethics Committee Protocol A2023/18 (Aplin), and under New South Wales Department of Planning and Environment Scientific License SL102790.

**Table 1:** Summary of age and sex of individually marked birds.

|  | Female | Male | Unknown | Total |
| --- | --- | --- | --- | --- |
| Adult | 25 | 25 | 0 | **50** |
| Juvenile | 19 | 3 | 9 | **31** |
| Unknown | 21 | 19 | 72 | **112** |
| Total | **65** | **47** | **81** | **193** |

**Table 2:** Summary of notation. Mathematical symbol, name, range of values (where applicable) and short description of relevant items in our model.

|  | Symbol | Range | Name | Description |
| --- | --- | --- | --- | --- |
| Parameters | $\alpha$ | $[0, \infty]$ | innovation rate | a baseline hazard that determined how frequently cockatoos innovated skills to defeat defences |
| | $\sigma_h, \sigma_c$ | $[0, \infty]$ | social learning probability | probability of social learning per timestep. |
| | $\lambda_h, \lambda_c$ | $[0, \infty]$ | decay factor | rate of information loss for households (motivation) and cockatoos (skill values). |
| | $\delta$ | $[0, \infty]$ | defence sensitivity | higher values meant that cockatoos more strongly preferred to attempt lesser defended bins. |
| Variables | $m_{i,t}$ | $[0, 4]$ | motivation level | Represents motivation to protect bins by household $i$ at time $t$. |
| | $D_{i,t}$ | $[0, 4]$ | defence level | Represents deployed defence level used by household $i$ at time $t$. |
| | $K_{i,t,s}$ | $[0, 1]$ | repertoire | Value cockatoo $i$ holds at time $t$ for skill $s$. $K = 0$ represents a naive state. |

#### 2.2.2 Transect observations of bin-opening

From July 1 to September 6, 2024, we conducted a total of 30 transects to observe SCC performing bin-opening, with transects conducted the evening prior to and the morning of rubbish collection. Two research assistants travelled random paths from random starting positions (both by foot and by car) throughout the study area and took note of any bin-opening attempts by SCC. Each transect began in a randomly assigned location as chosen from a grid of 22 blocks that measured 200×200 meters. Evening transects lasted for approximately two hours, from 16:00 until sunset. Morning transects lasted for approximately three hours, from 6:00 until 9:00. Transects did not have a pre-defined path, but the geographic area was small enough that research assistants could sample the entire area at least once within a transect if by foot, and several times if by car. Combined with the random starting point, this provided the best practical chance at collecting a random sample of behaviours. For each observation we recorded the location, time, identities of birds who interacted with the bins, the type of protection used on the bin, and what stage of the opening process the openers achieved. Opening stages 1-5 are defined and detailed in Klump et al. (32), and are summarized as stage 1) prying the lid upwards with the bill, 2) raising the lid into an open position, 3) holding the lid open with foot while repositioning the body for walking, 4) walking towards the hinges such that the lid opens further and 5) flipping the lid over entirely. A protection was considered as defeated if either the bird went on to fully open the bin (stage 5), or if the bird was able to reach stage 4 of the opening and extract food. We also recorded 112 observations of unmarked birds interacting bins, although these were excluded from analysis since we intended to characterize the knowledge states of identifiable birds in the present study. We note that we also habituated and marked birds in 2023, but do not present behavioural data from 2023 in the present manuscript, as we did not conduct a knowledge assay in 2023.

#### 2.2.3 Knowledge assay

To supplement the above dataset, we provided SCC access to bins at the habituation site with or without protections. Assays were conducted over nine non-consecutive days (August 8-23, 2024) for 90-180 minutes per day. We provided three bins arranged in a triangle with 2m of separation to reduce social interference during attempted openings. We provided unprotected bins for 3 days, bins with a loose weight on top for two days, bins with an empty water bottle placed in the hinge for two days, and bins with a purpose-made lock for two days. These were selected as representing different classes of protection, of the types most commonly observed. For each attempt at opening a bin, we recorded the identity of the bird, whether or not the defence was defeated (for defended bins), what stage of bin opening the attempting SCC reached (according to the ethogram defined by Klump et al. (32)), and whether or not there was social interference by other individuals. Every 30 minutes, we recorded all birds present to determine which birds were present but did not attempt to open the bins. We excluded 4 observations of unmarked birds.

This assay was conducted during the same time-period that bin-opening assays were being conducted. However, it ultimately would have little impact on the overall uptake rate of the population. Firstly, SCC are exposed to bins year round. The few hours comprising the assay was very insignificant exposure time compared to the overall exposure that occurs naturally. Furthermore, we believe that it takes months of exposure for bin-opening behaviour to be acquired, as it is a complicated 5-step action that requires adequate strength (32). We find only a minority of birds successfully opening bins.

### 2.3 Agent based model

#### 2.3.1 Overview of schedule

In order to better understand the dynamics of behavioral arms races, we created a spatially explicit simulation model of two competing populations of leaky accumulator learning agents, representing households and SCC. In each timestep (*t*), representing a bin-collection day, households decided whether they would leave their bin unprotected, or to protect their bin using a maximum of three defences. Their choices depended on a motivation level, which was affected by past experience and by the motivation levels of neighbors, according to a social learning rate (*σ_h_*).

SCC then moved to a household following a movement rule, and attempted to use skills from their repertoire *K* to open the bins. Four different skills directly corresponded to the defense levels, and could either be individually learned with a constant innovation rate (*α*), or learned from observing others perform successful actions according to a social learning rate (*σ_c_*). SCC were only able to open bins if they held the corresponding skills in repertoire (e.g. to open a D2 bin, SCC must perform D2, D1 and D0). If a SCC agent successfully opened a bin, the actions used were reinforced. Otherwise, their values decayed over time, causing them to potentially forget actions. The precise rate of reinforcement and decay are defined as reasonable values in this simulation, and we took steps to validate the relationship between the two detailed in Section 3.4, but would need to be verified at some point with the appropriate empirical data.

#### 2.3.2 Household agents

We defined households as cells in a grid, where each node represented a single household with one bin. Households were characterized by a defence level *D_t_* ∈ [0, 3], where 0 represented unprotected bins, and 3 was a maximal defence level unable to be defeated. Each household was modelled as a leaky accumulator, keeping track a motivation value (*m_i,t_*; initialized as *m_i,_*_0_ = 0), which was updated at each timestep depending on the outcome of its defensive effort and a natural decay over time:

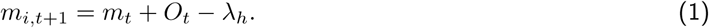

Outcome *O_t_*could increment motivation depending on whether a bin was opened, where

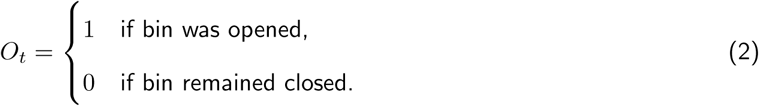

As evidenced by our empirical data, protections carried some cost in terms of effort to deploy, represented by the decay factor *λ_h_*. For simplicity, we assumed that all defences were equally costly. We constrained 0 ≤ *m* ≤ 4, with households choosing *D_t_* based on whether *m_t_* fell between defined threshold levels *T* = [1, 2, 3]:

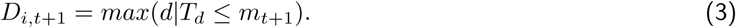

For example, if 0 ≤ *m_i,t_ <* 1 the bin was undefended, and if 1 ≤ *m_i,t_ <* 2 the household chose *D_i,t_*= 1. Setting *O_t_* = 1 for unsuccessful defences meant that a household would increase defence levels immediately after a successful opening. While this is not entirely realistic, it allowed us to easily compare decay rates between households and SCC. Furthermore, we found relatively few cases where households jumped from no protections to multi-protections from 2023 to 2024, suggesting that step-wise increases in protection-level are realistic (Figure S2).

Households could potentially be influenced by the decisions of their four cardinal neighbors according to a social learning probability (*σ_h_*). If an agent socially learned, they adopted the mean motivation of their neighbors.

#### 2.3.3 SCC agents

SCC agents were modelled as leaky accumulators, accumulating values for four skills *s* in repertoire *K*. We chose to model values for each skill separately (c.f. singular *m* value for households) because our empirical data showed that it is possible to know how to defeat protections without necessarily knowing how to open bins. After attempting to open bins, SCC updated their values *K* using

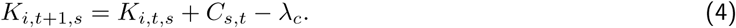

Skill values were reinforced using

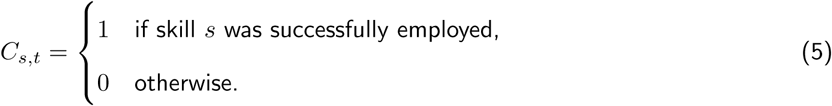

Values were constrained such that 0 ≤ *K_s_* ≤ 1. An individual could only perform skill *s* when *K_s_ >* 0, meaning that skills could be forgotten over time according to the decay factor *λ_c_*. When more than one SCC was present at a household, and a bin was successfully opened by bird *j* using skill *s*, any SCC naive to skill *s* could potentially acquire it by copying the demonstrator’s value *K_j,t,s_* according to social learning probability *σ_c_*. This meant that birds could acquire the skill *s* depending on a draw from a Bernoulli trial where *p*(socially learn) = *σ_c_*.

SCC necessarily had to choose which bin to attempt each timestep. SCC have a homerange of 2-3 square km (41; 42), and it is reasonable to assume that they are generally unconstrained in choosing which bins to approach. We tested two movement rules: random movement, where all households had an equal chance of being chosen, and defence sensitive movement, where SCC preferentially chose houses with lower defence levels. Given vector *D* of household defense levels, movement probabilities were based on the relative “unprotectedness” (*u*) of each household, calculated as:

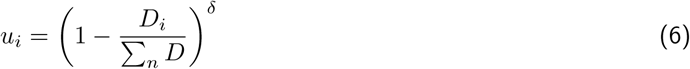

where *δ* controlled sensitivity to differences in unprotectedness. We found that *δ* = 500 let agents prefer lower-defended households, but still allowed for some randomness in their choices. The probability of an agent moving to a given household was given by:

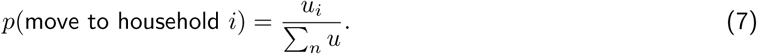

If all defense values were *D*0, agents moved at random across all locations.

#### 2.3.4 Measurements

At each timestep, we measured the following metrics:

1. the sum of defence levels of all households as a measure of overall cost to the human population.
2. the mean, lower 2.5 quantile and upper 97.5 quantile of household motivation levels.
3. The number of bins that were successfully opened.
4. The number of SCC that knew each skill (*D*0 − *D*3).

At each timestep, we calculated the cost/benefit ratio for households (cost per bin protected) as the cumulative sum of defences used divided by the cumulative sum of opened bins. Using a cumulative sum accounted for the cost of time spent maintaining defences, and we assumed a higher cost for higher levels of defence.

#### 2.3.5 Scenarios

For each of the scenarios below, we conducted 100 simulations that each ran for 1500 timesteps.

1. **vs.1**: One household and one SCC, such that effects of movement and social learning are removed.

We used this simple scenario to understand how different innovation rates, decay rates, and efficacy regimes affected simulation dynamics, and used these results to fine-tune parameter settings for the remaining scenarios. Results for this scenario are fully presented in supplementary text S1. We then increased the population sizes to *N* = 100 households and *N* = 20 SCC and tested different combinations of learning strategies:

2. **Household IL, Cockatoo IL**: Both households and SCC only individually learn.
3. **Household IL, Cockatoo SL**: SCC may socially learn skills.
4. **Household SL, Cockatoo IL**: Households may socially learn their neighbors’ motivation level/defence level.
5. **Household SL, Cockatoo SL**: Household and SCC may socially learn.

Animations of exemplar simulations for the above scenarios are shown in Video S1-S4. Given that we have evidence that both humans and SCC socially learn, we allow for this in the remaining scenarios where we then explored the outcomes of potential responses by councils. The grid world was divided into two equally sized councils (A and B) of 50 households each. Households from council A could not socially learn from council B. We then tested the following coordinated responses:

6. **Universal policy**: After 500 timesteps, both councils A and B implemented mandatory *D*3 protection for all households. The policy ended after 1000 timesteps.
7. **Unilateral policy**: After 500 timesteps, council A implemented mandatory *D*3 protection for all households. Council B did not implement a protection. The policy ended after 1000 timesteps.
8. **Pre-emptive unilateral policy**: At the first timestep, council A implemented mandatory *D*3 protection for all households. Council B did not implement a protection. The policy ended after 500 timesteps.

Animations of exemplar simulations for the above scenarios are shown in Video S5-S7 for random movement, and Video S8-S10 for defence-sensitive movement. We set the policy periods to be equal to each other so that the results could be comparable. While policies could be permanent in reality, we continued to measure the system after the policies ended to evaluate the system dynamics after the policies’ end. The end of policies could also be interpreted as a scenario where the defence degrades over time, as we observed many broken protections that were not replaced.

## 3 Results

### 3.1 Bin protections

We recorded protections of 1419 bins at 643 unique addresses in the study area over two years (386 addresses and 732 bins in 2023, 453 addresses and 687 bins in 2024), with 196 addresses recorded in both years. This represented approximately 75% of the local population of 858 dwellings. Overall, 852 bins were undefended (no defensive measure, or a broken defensive measure), 517 bins had a single defence (e.g. a single lock or bottle), and 50 bins had multiple protections with either several of the same defence (e.g. two loose weights) or a combination of protections (e.g. a lock and bottle). We observed a trend for decreasing protection levels, as the percent of unprotected bins in our survey increased from 55.6% to 64.8% between 2023 and 2024 (up from 50.7% in 2019 (36)). Amongst bins with single protections, locks accounted for 36% of all protections, followed by 17% with weights, 16% with sticks, and 13% with bottles. In Stanwell Park, locks were installed by homeowners, although in 2023 the suburb of Campbelltown provided locks to homeowners in a trial scheme (43). Other protection types that accounted for less than 10% of observations included ropes, springs, hooks, wires, bungees, and strategic blocking of the bin-lid by the placement of bins (see Table S1 for details). When aggregating observations by address and taking the highest protection observed, we found that in general, residents decreased their protection levels from 2023 to 2024 (Figure 1A), also matching the trend found at a bin-level analysis (Table S2). The proportion of households with no protections observed rose from 38% to 51%. The proportion of addresses with single protections fell from 55% to 45%, and multiple protections fell from 7% to 3%. We found that broken protections only comprised 4% of all unprotected bins in 2023 and 3.3% in 2024, suggesting that overall decreases in protections were active decisions, rather than a more passive process of not replacing broken protections.

**Figure 1:**
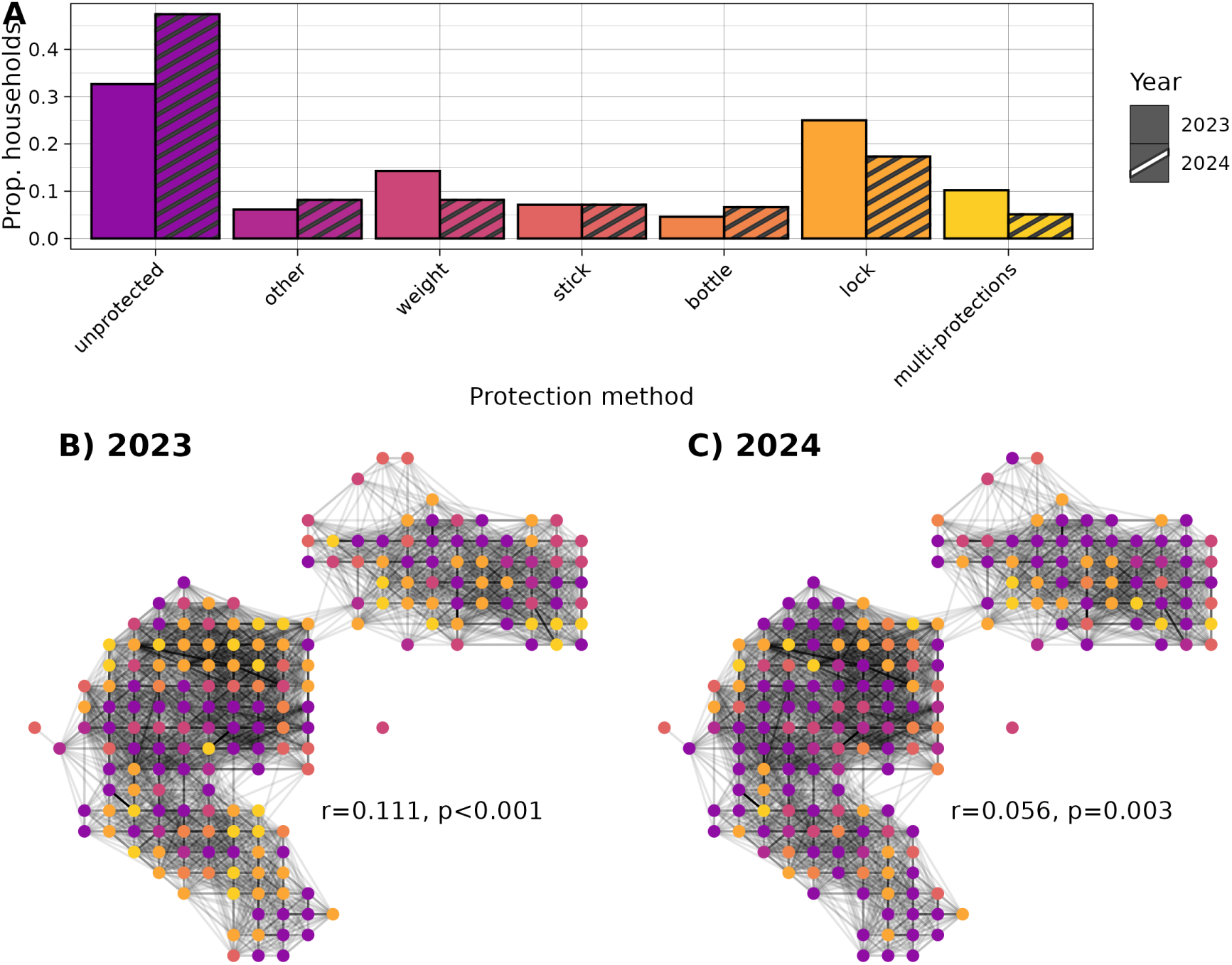
Household-level changes in protection methods. A) Proportion of unique addresses using each protection method. Overall protection levels decreased from 2023 to 2024. B, C) Nodes are unique addresses colored by protection methods, edges are shaded by inverse driving-distance. Connections that are *>* 200m are removed for visualization. Protection methods were positively spatially assorted in 2023 and 2024.

The binary decision to protect bins, and the efficacy of bin protections were observed to be significantly positively assorted by geodesic and driving distance in 2023, and non-significantly positively assorted in 2024 (Table S3). However, in both years, protection method was significantly assorted as measured by both geodesic and driving distance (Figure 1B, C).

### 3.2 SCC knowledge states

Over the course of 18 bin-opening transects, we record a total 205 observations, 87 of which were of a marked bird and where the entire sequence of behaviour was recorded. These 87 attempts of bin opening were by 31 marked individuals on 43 unprotected bins and 44 protected bins (Table S4). Only about 30% of these attempts were successful, with 25 successful openings by 11 marked cockatoos. 20 of these successful openings were on unprotected bins. It was also possible that birds could be knowledgeable about how to defeat a protection, but not yet know how to open a bin. In this case, a single bird was also observed defeating a loose weight but failed to fully open the bin. We also observed several failed attempts at defeating stick, spring, rope and fixed-weight protections.

In addition to this observational data, we conducted a controlled knowledge assay where we presented birds with bins equipped with commonly found protections of increasing efficacy (Figure 2A-D). Over 9 days of data collection, a total of 153 marked birds visited the assay site, and we observed a total of 301 attempts to open bins by 48 marked birds (Table S5). Most birds attempted to open unprotected bins, followed by bins with a lock, bins with a bottle, and bins with a loose weight. Success in defeating the protection and opening the bin decreased with protection efficacy. When looking at an individual level (summarized in Table S6), 3 birds were observed as knowing how to defeat two protections, 11 individuals were observed to defeat one protection. We recorded 8 individuals who could defeat protections, but did not successfully open the bin afterwards.

**Figure 2:**
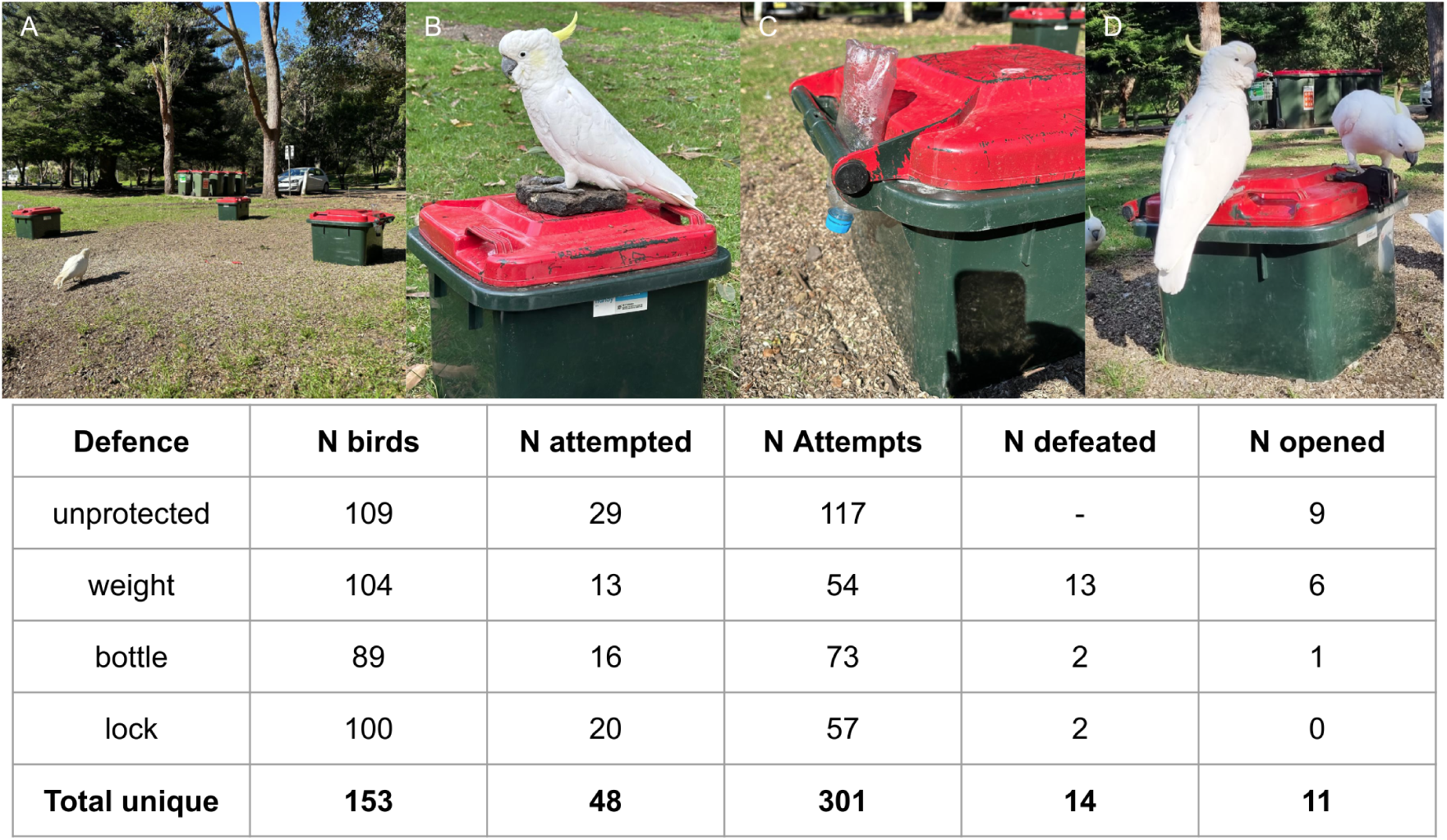
Summary of knowledge assay. A) Configuration of bins. B) Bin with loose weight. C) Bin with bottle in hinge. D) Bin with lock. In table, columns include the protection type, the number of birds exposed to the task for longer than 30 minutes, the total number of attempts, the number of unique birds to attempt, the number of birds that successfully defeated the defence, and the number of birds who then went on to open the bin. In parentheses is the number of unique birds who were observed opening a bin independent of protection level.

When combining data from transects and the assay, we had data for 154 of the 193 marked SCC. We identified 22 SCC with some skill at bin-opening (summarized in Table S7). 17 SCC were confirmed to fully know how to open bins (7 adult males and 2 males of unknown age, 1 adult female, 2 juveniles and 5 SCC with unknown age or sex). Only 5 of these birds were observed opening in both the assay and the wild, highlighting the value of combining these two datasets given how difficult it is to observe the naturally occurring behaviour. 13 birds could defeat weights, only 3 could defeat bottles, and surprisingly, 4 could defeat locks (see Video S11 for an example from the wild). During transects, we additionally observed 1 bird defeating block protection. A further 49 marked birds were observed unsuccessfully attempting to either open bins or defeat protections.

### 3.3 Arms race model

The data above suggest several important considerations for our model. Bin-opening has been observed in this locality for over 10 years, yet we identified relatively few birds as able to open bins, representing about 11% of marked residents about which we had observational data. Supporting previous work, we found that 10 of 11 bin-openers of known sex were male. Across the years of observation, 42 − 52% of DNA-sampled individuals were male, suggesting a fairly even population sex ratio. This further suggests that bin-opening behaviour is almost entirely restricted to males. Yet even given this sex-bias, the low percentage of learners means we can conclude that the innovation rate and social learning rate must be relatively low. Given the level of turnover in bin protections by households, we know that this is a dynamic process where motivation for using protection can change over time. Given the shift towards unprotected bins and de-escalations from multiple protections, we can assume that these strategies might be costly to maintain. We constructed our model to be able to explore these considerations, and the consequences for system dynamics, as well as how the system might respond to bottom-up versus top-down strategies for mitigating this human-wildlife conflict.

### 3.4 Individual and social learning

We first explored the simplest case where *N_h_* = 1 household and *N_c_* = 1 SCC, such that social learning was *de facto* impossible, and the effects of movement rules were removed. The dynamics of this condition lead to identification of several critical relationships in our model that determine its dynamics. We briefly describe these below, although these points are illustrated by exemplar simulation runs in supplementary text S1, figures S3, S4, and S5. Higher innovation rates (*α*) reduced the time to achieve maximal protection levels. In reality, innovation must be relatively rare compared to social transmission, as bin-opening behaviors are recent and geographically limited. Behavioral extinction was influenced by the decay rates (*λ_c_*, *λ_h_*). If humans de-escalate faster than SCC forget, escalations to higher protection levels can occur; otherwise, if *λ_c_*is high relative to *α* and *λ_h_*, SCC forget the behavior rapidly, limiting full escalation to *D*3. The efficacy of protection levels also shapes outcomes. Assuming SCC can reliably bypass protections leads to full escalation, while increasing defense efficacy reduces success rates and creates oscillations or unpredictable dynamics. Given the observations above, for the following simulations we have set innovation rate *α* = 0.005, *λ_h_* = 0.05, *λ_c_* = 0.01 (such that motivation decays faster than SCC memory), and we assume increasing efficacy of defences *p*(defeat|know skill) = [1.0, 0.5, 0.25, 0.125].

We increased the number of households to *N_h_* = 100, and the number of SCC to *N_c_* = 20. This necessarily introduced effects of movement rules of SCC, and we tested both “random movement” (Figure S6), and “defence sensitive movement” (Figure S7). We compared different combinations of whether or not households or SCC could socially learn about their neighbors’ motivation level or their associates’ skills respectively. SCC social learning increased the spreading rate of skills once innovated, as expected. Interestingly, for either movement rule, when households used social information they reduced the relative costs compared to if they acted independently (Figure 3).

**Figure 3:**
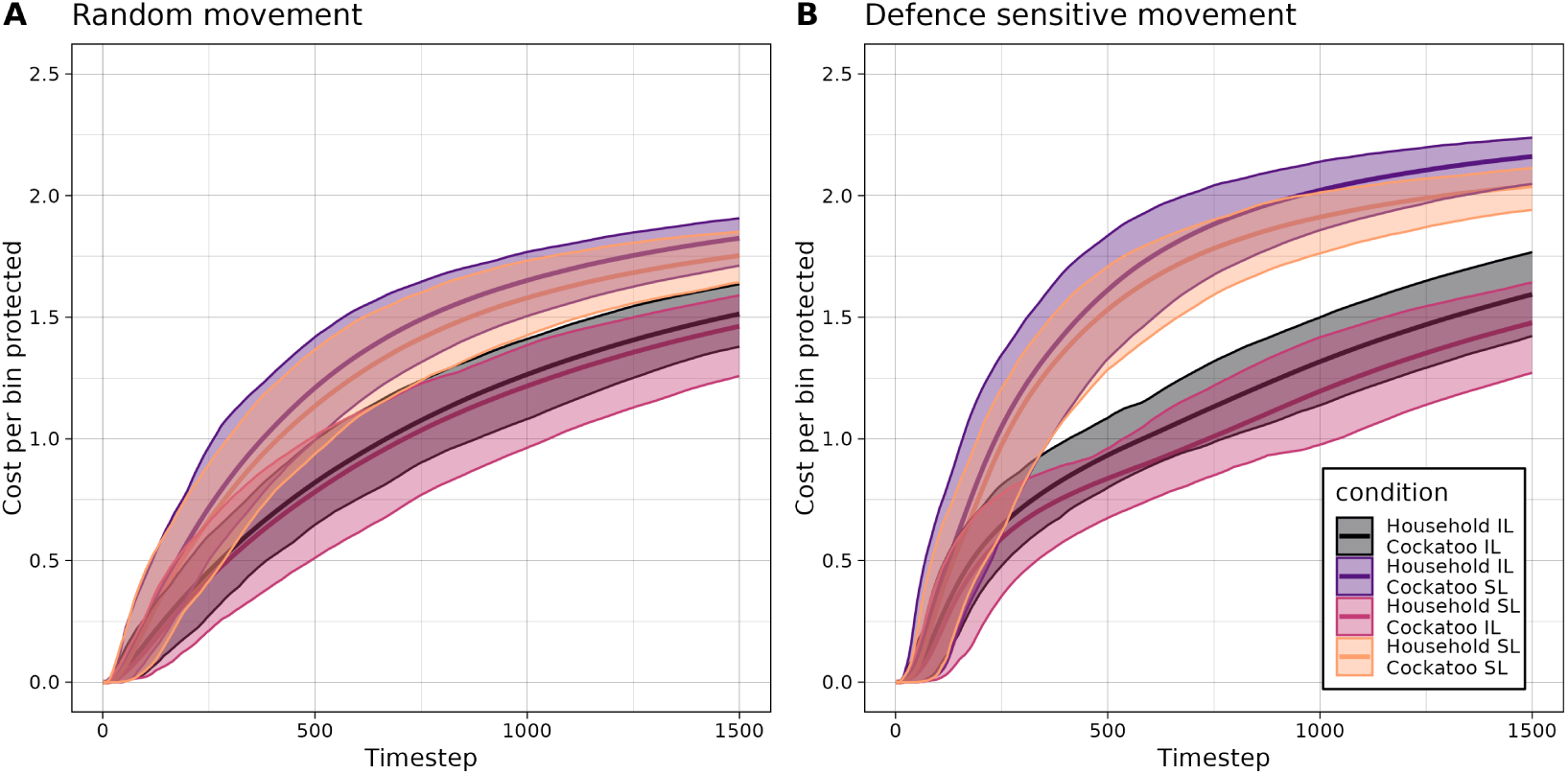
Social learning can affect cumulative costs per benefit. Mean (95% interval shaded) cost per protected bin over time, colored by condition. Highest costs arise when sulphur-crested cockatoos socially learn and households do not. If households use social learning, they can reduce the collective costs of protection, regardless of whether cockatoos are socially learning.

We observed that independent of how agents learned, defence sensitive movement greatly reduced the possibility of full escalation to *D*3, as SCC avoided higher defensive households (Figure S6 c.f. Figure S7, panels C,F,I,L). This resulted in 1) households de-escalating after introducing higher protections due to lack of visitations by SCC, and 2) SCC not learning how to defeat higher defences because their overall exposure was lower.

### 3.5 Coordination of defences

We wanted to understand the consequences of a top-down coordinated policy, compared against *laissez-faire* protection, where households make decisions independently. In the following simulations, we divide households into two distinct neighboring councils, and assumed that both households and SCC can socially learn (although households only learned from those in their council). We compared different policy scenarios to a baseline scenario with no policy from either council (Figure 4A, B, C), whose simulation dynamics were identical to the SL/SL case above. We tested these scenarios with both SCC random movement (Figure S8) and defense-sensitive movement (Figure 4).

**Figure 4:**
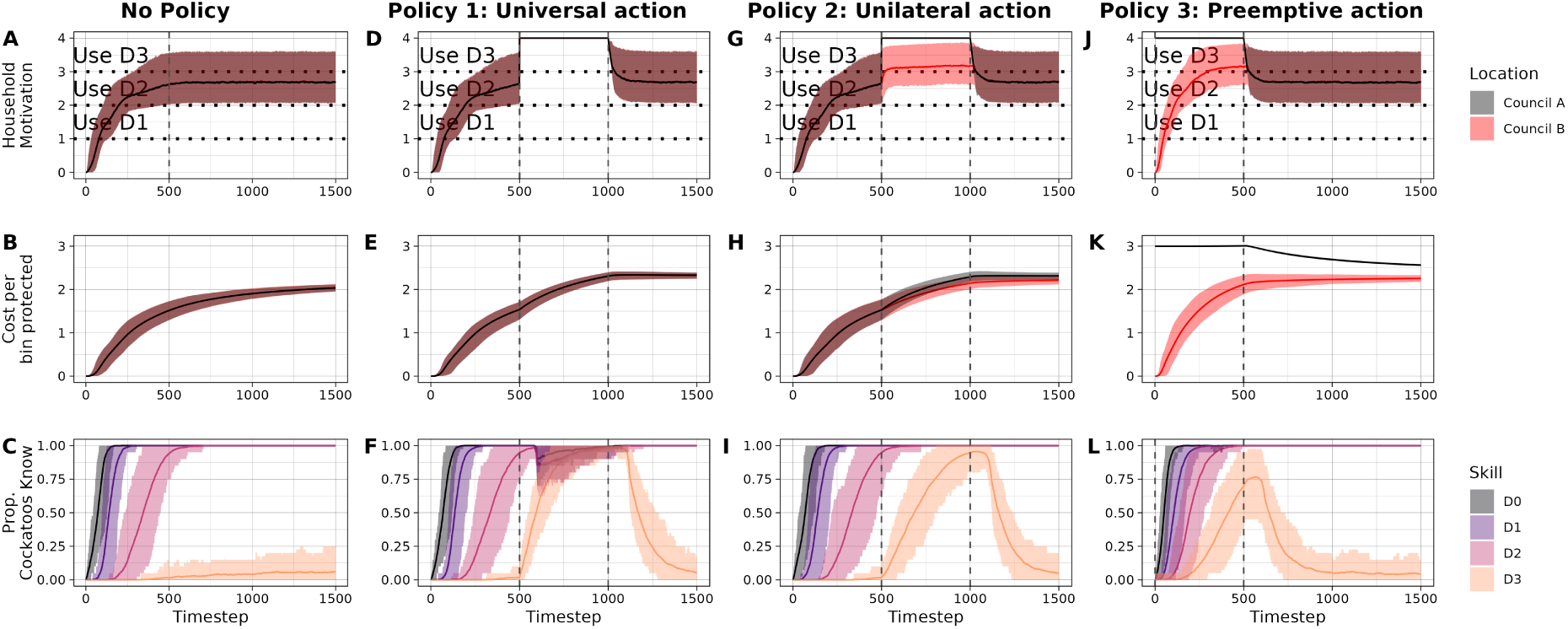
Coordinated policies can have unintended consequences for neighboring areas. Average values and 95% intervals (shaded) under four different policies (columns), for two neighboring councils (black: policy zone, red: neighboring area). Policy start and end points are indicated by dashed vertical lines. Compared with no policy (A, B, C), enforcement of *D*3 defences either universally (D, E, F), unilaterally (G, H, I) or preemptively (J, K, L) causes a sharp increase in the number of cockatoos with larger repertoires, because it necessarily provides them with exposure to all three defences. This raises the average household motivation (D,G,J) and overall cost per bin protected in both councils (E, H, K).

A universal action in which all households escalated to *D*3 after 500 timesteps and maintained the defence for 500 timesteps, resulted in an immediate decrease in the number of successfully opened bins (Figure S9B,F). Importantly, this did not result in behavioral extinction, as it was possible (although improbable) for a bin to be opened. In fact, maximal exposure to *D*3 defences meant that there were more opportunities to innovate or socially learn the skills to defeat *D*3, and the proportion of SCC with *D*3 skills sharply increased compared to the baseline with no policy (Figure 4C,F).

We then tested a unilateral action where Council A implemented *D*3 while Council B did not implement any policy. If SCC had defence sensitive movement, implementing a unilateral policy displaced SCC into Council B. This resulted in a spike in successfully opened bins in Council B (Figure S9G), at least until they adapted their defence levels to the influx of birds by elevating it above what it would have been otherwise (Figure 4G). The cost per protected bin for both councils also rose higher than in the no-policy scenario (Figure 4H). This scenario also resulted in hastened learning of the skill to defeat *D*3 (Figure 4I). A pre-emptive, unilateral action also elevated household motivation in Council B higher than it would be otherwise (Figure 4J), and skills to defeat *D*3 were innovated earlier and spread faster than in other scenarios (Figure 4L). Importantly, when SCC movement was entirely random we found no differences in the cost benefit ratio of Council B between the no policy scenario and unilateral scenarios (Figure S8 B, H, K). However, when SCC preferentially avoided protected bins, the cost of protecting bins was elevated under policies 2 and 3, despite council B not enacting any mandatory protections (Figure 4 B, H, K).

## 4 Discussion

By combining observational, experimental and theoretical simulation approaches, we have found new insights into the potential innovation arms race occurring between humans and sulphur-crested cockatoos in urban areas of eastern Australia. First, we added an additional two years of data to Klump et al (36), finding that the method and efficacy of bin protection measures used by local residents in our study area in 2023 and 2024 are significantly spatially assorted. This supports the claim that bin protections are socially learned, suggesting that people are exchanging information with their neighbours and copying their protection methods. Despite widespread concern about bin-opening, our data show a decrease in the use of bin protections between 2023 and 2024, also slightly lower than in 2019 (36). This decline suggests that protection measures may be costly or inconvenient to maintain, leading to de-escalation over time. We found that a small minority of unprotected bins were due to broken protections that residents had not replaced, suggesting that most de-escalations were active decisions to remove protections, rather than failing to maintain them. In some cases, residents may also be adapting their behavior by tolerating the presence of cockatoos or modifying their behavior in ways not captured by our study (e.g. using strategic timing to bring bins out immediately before collection).

Our behavioral observations of cockatoos suggest that relatively few birds have mastered bin-opening or protections, although the number of bin-openers (17 total) was more than the 9 knowledgeable birds identified in the same population in 2019 (32). This might indicate that more SCC have learned to open bins since the first study, although 2019 estimate might also be low, as there was no knowledge assay to complement bin-opening transects. Our controlled knowledge assay confirmed that some individuals can defeat specific protections but may not necessarily know how to open bins (or at least didn’t demonstrate it), leading to variation in skill levels within the population. Given that SCC who know how to fully open a bin are relatively rare, yet bin-opening behavior has persisted in this suburb for nearly a decade suggests that the innovation rate is likely low, with social learning playing a key role in skill acquisition. We identified a number of birds unsuccessfully attempting to open bins or defeat protections (N=49, more than N=27 in 2019), that may be actively learning, and future longitudinal studies of the progression of the behavior are needed to better understand what precisely is individually and socially learned by the cockatoos. It could be the case that certain components are socially learned, while the remainder of the behavioral sequence is individually learned, as found in great tits during an experiment with a two-action puzzle (44).

Parallels can be drawn between this behavioral “arms-race” typified by this bin-opening nuisance conflict and the more general case of evolutionary arms races. One key concept of evolutionary arms races is the “Red Queen Hypothesis”, which describes co-evolutionary dynamics where antagonistic species must continuously adapt to maintain their relative fitness (45). Similarly, residents and SCC continually update their strategies, with each side locked in an adaptive race potentially driven by cultural transmission rather than genetic evolution. Our findings highlight that social learning accelerates adaptation in both humans and SCC, preventing either from easily outpacing the other. Additionally, we find that policy interventions, such as bin-locking programs, can inadvertently drive innovation in cockatoos, analogous to how increased selection pressure in natural systems can spur rapid evolutionary responses in prey or pathogens. The Red Queen Hypothesis has also led to the question of why species do not adapt indefinitely, (i.e. why do we ever observe extinction), and what mechanisms limit infinite adaptation? One answer is the associated cost of adaptation (46). For humans, the cost of bin protection arises both from the money spent paying for protections such as locks, but also the time invested in setting up and maintaining protections. For cockatoos, the cost appears to largely be in the time invested in learning to open-bins and defeat protections. Furthermore, the weight of bin-lids might prevent many SCC from ever being able to acquire the behavior, and go some way to explain the male bias in the behavior.

Our simulations emphasizes some critical points for the human response to bin opening. Decision-making that includes some information about what one’s neighbors are doing was found to mitigate the overall costs of bin protection. Coordinated responses in one area (e.g. introduction of council-wide locks) reduce the opening rate, but also could potentially push already knowledgeable birds into novel areas. This will increase the bin-opening rates in these novel areas, at least temporarily until residents respond, and raises their cumulative costs higher than if no policy had been enacted. Furthermore, such policies can encourage the innovation and spread of skills to defeat novel protections by increasing birds exposure rates to the task, assuming that the protections are not perfectly effective. While the scale of our model’s population was too small to show this, we can also predict could spread bin-opening to birds that would otherwise be un-exposed to the behavior. Finally, even if we assume that a defence has perfect efficacy, as soon as the policy ends, any underlying innovation rate ensures the bin-opening behavior will eventually be re-innovated. We acknowledge that our simulation model was greatly simplified for tractability. More research is needed to understand the sensitivity of households to changes in defence levels after successful opening and defending episodes. Also, cockatoos’ movement is likely not independent of each other, and it is still unclear how cockatoos make choices of which bins to interact with. Individuals might develop habitual routes (42), or choose bins which were previously rewarding, or perhaps are attracted to other cues, such as the smell of food waste or the presence of others. Introducing socially influenced movement rules fell outside the aims of the current manuscript, but also are likely change model dynamics.

In summary, our study highlights how human-wildlife conflict is not a static problem but a dynamic system, where interventions must consider long-term feedback loops. A single fix, such as providing all households with locks, may not halt the escalation. We find evidence to support that bin-protection choices are likely socially influenced, and that the bin-opening behavior is continuing to spread in SCC. We encourage social scientists to explore the human dimension of the bin-opening phenomenon to further explore when and why residents decide to protect, escalate or de-escalate their protections. Finally, we illuminate how policy choices might drive innovation in SCC and inadvertently escalate the arms-race.

## 5 Data Availability

Supplementary videos, as well as code and data for the agent based model, statistical analyses, and main text figures are publicly available on Edmond at https://doi.org/10.17617/3.XD6UEW (47).

## 6 Acknowledgements

We thank the field assistants involved in collecting data for this project: Lakesha Smith, Christophe Turcotte-van de Rydt, Thea Osmond, and Anna Thieben. We are very grateful to the local residents of Stanwell Park and Sutherland Shire who welcomed us to work in their communities. We thank Diversity Arrays Technology for performing the genetic analysis, and to Kyle Ewart for giving advice on this analysis. We additionally thank Nitzan Shahar, Arnault-Quentin Vermillet, Britt Anderson, Dimitris Askitis and Kyveli Kompatsiari for their contributions to the conceptualization of the agent based model at the 2024 Lorentz Center Workshop on Cognitive Modeling of Complex Behavior in Leiden, NL. This work was supported by the Cultural Evolution Society Transformation Fund, funded by the John Templeton Foundation, Grant 61913. The opinions expressed in this publication are those of the author(s) and do not necessarily reflect the views of the John Templeton Foundation. MC and LMA were supported by the Swiss State Secretariat for Education, Research and Innovation (SERI) under contract number MB22.00056. MC had additional support from the Centre for the Advanced Study of Collective Behaviour, funded by the Deutsche Forschungsgemeinschaft (DFG) under Germany’s Excellence Strategy (EXC 2117-422037984). During the final stages of this project, BCK has been funded by the Vienna Science and Technology Fund (WWTF) [10.47379/VRG21011]. All work was conducted under AEC Ethics protocol A2023/18 and NSW Scientific License SL102790.

## 7 Author Contributions

Conceptualisation, M.C., E.D., B.C.K., L.M.A.;

Methodology, M.C., E.D., B.C.K., L.M.A,;

Software, M.C., E.D.;

Investigation, M.C., E.D.;

Visualization, M.C.;

Writing - Original Draft, M.C.;

Writing - Review & Editing, M.C., E.D, B.C.K., L.M.A..

