## supplementary for "Temporal dynamics and policy implications in an innovation arms race"

### Supplementary Materials

#### Text

- **Text S1**: Model dynamics under 1 versus 1 scenario.

#### Figures

- **Figure S1**: Example photographs of common bin defences.
- **Figure S2**: Year over year changes in protection type (unprotected, single protection, multiple protections).
- **Figure S3**: Effects of innovation rate (1 household vs. 1 cockatoo).
- **Figure S4**: Effects of decay rates (1 household vs. 1 cockatoo).
- **Figure S5**: Effects of defence efficacy (1 household vs. 1 cockatoo).
- **Figure S6**: Simulation dynamics of different learning types under random movement.
- **Figure S7**: Simulation dynamics of different learning types under defence sensitive movement.

#### Tables

- **Table S1**: Proportion of addresses using each protection method.
- **Table S2**: Summary of each protection type by year.
- **Table S3**: Bin protection assortativity values by year.
- **Table S4**: Individual-level summary of bin-opening transects.
- **Table S5**: Summary of knowledge assay
- **Table S6**: Individual-level summary of knowledge assay.
- **Table S7**: Individual-level summary of assay and transect data combined.

#### Videos

- Video S1: Exemplar simulation: household IL / cockatoo IL, random movement
- Video S2: Exemplar simulation: household IL / cockatoo SL, random movement
- Video S3: Exemplar simulation: household SL / cockatoo IL, random movement
- Video S4: Exemplar simulation: household SL / cockatoo SL, random movement (same as policy 0)
- Video S5: Exemplar simulation: Policy 1, random movement
- Video S6: Exemplar simulation: Policy 2, random movement
- Video S7: Exemplar simulation: Policy 3, random movement
- Video S8: Exemplar simulation: Policy 1, defence movement
- Video S9: Exemplar simulation: Policy 2, defence movement
- Video S10: Exemplar simulation: Policy 3, defence movement
- Video S11: Cockatoo ORG observed defeating a lock protection in the wild.

### S1 Model dynamics under 1 versus 1 scenario

Cockatoo innovation rate ($\alpha$) was directly related the frequency of increased threat to bins. Thus, increasing $\alpha$ decreased the time it took to reach maximal protection levels (Figure S3). Our data suggests that the true innovation rate is quite low relative to the social transmission rate due to the recency of bin-opening and the lack of reports from other major cities with cockatoo populations, despite bins being a constant feature of residential communities for decades.

The relationship between decay rates for households and humans ($\lambda_h$, $\lambda_c$) largely determined behavioral extinction. If $\lambda_c > \lambda_h$, protections are used for longer than the cockatoos can keep skills in repertoire (Figure S4 A,B,C). Once a skill is forgotten, it may only be regained through innovation, determined by $\alpha$. If innovation is low, and cockatoos are forgetful, there will be infrequent periods where the behavior is innovated, an consequential increase to $D1$. However, the cockatoo quickly forgets the skill, and the household returns to $D0$. An innovation to $D1$ is unlikely to occur before this de-escalation. If $\lambda_c < \lambda_h$, humans will de-escalate before cockatoos have forgotten the behavior, leading to more successful attempts, and an eventual escalation to $D3$ (Figure S4 D,E,F). Given that bin-opening continues to be a problem, and that we have observed birds to be able to defeat protections, we conclude that cockatoos are not so forgetful. Given the number of de-escalations we observed in protections used after 1 year, we conclude that it's likely that $\lambda_h > \lambda_c$.

The efficacy of each defence level also affected simulation dynamics by modulating the probability of successful skill usage by cockatoo. One could make several types of assumptions about efficacy. The simplest is to assume that a cockatoo can reliably defeat any protection once it has innovated a solution (e.g. $p(\text{defeat}) = [1.0, 1.0, 1.0, 1.0]$). This led to an escalation to $D3$ as the cockatoo sequentially innovated all 4 skills, with no opportunity for de-escalations, and a 100% success rate for the cockatoo (Figure S5 A,B,C). One could also assume a threshold scenario, where $D3$ represents an indefatigable (although still costly) protection, such as locks had been considered prior to this study (e.g. $p(\text{defeat}) = [1.0, 1.0, 1.0, 0.0]$). This resulted in a final oscillating state of defences between $D2$ and $D3$ (Figure S5 D,E,F). After applying $D3$, the household loses motivation until it reverted to $D2$, which the cockatoo defeats, causing the household to reapply $D3$. Finally, one could assume that there is some increasing efficacy to defence levels, such that the probability of defeating each successive defence decreases (e.g. $p(\text{defeat}) = [1.0, 0.5, 0.25, 0.125]$). This created less predictable fluctuations in motivation levels and cockatoo's value of defence skills, and allowed for the possibility of behavioral extinction, although this would depend strongly on decay rates. If the interval between successful openings was still shorter than it took for cockatoos to forget their skills, then a similar escalation pattern as before is observed (Figure S5 G,H,I).

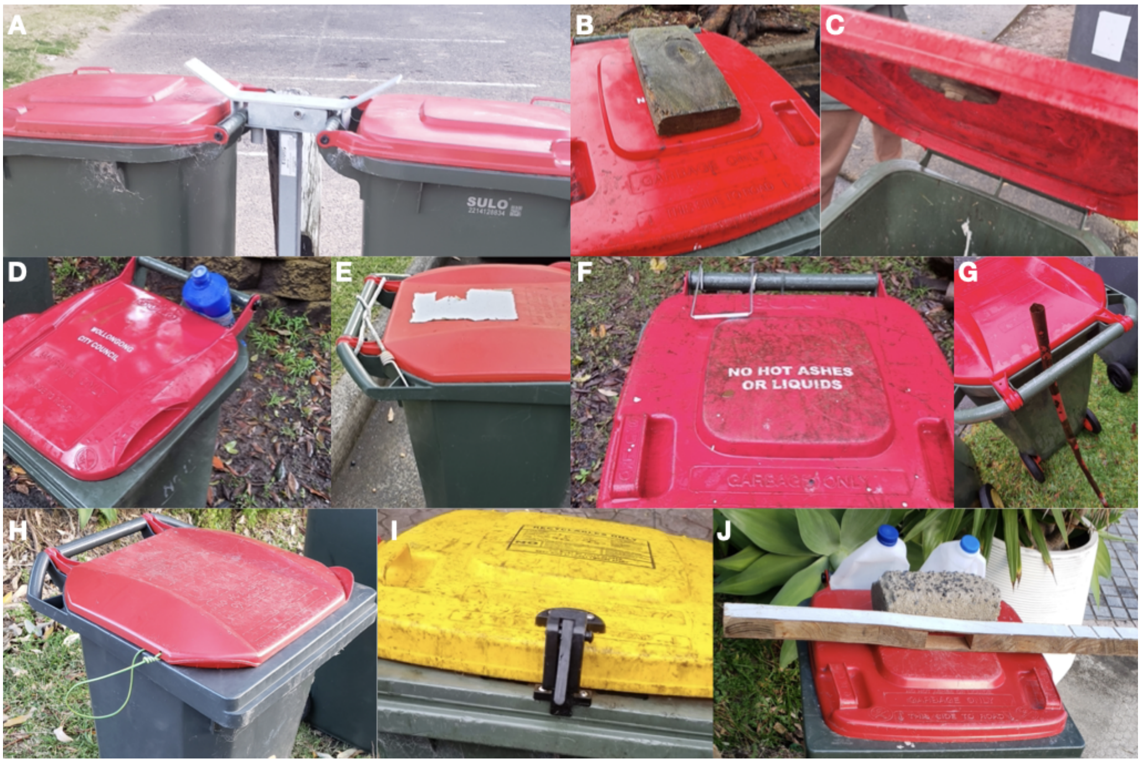


Figure S1: Example photographs of common bin defences. A) Block, B) loose weight, C) fixed weight, D) bottle, E) bungee-cord, F) spring, G) stick, H) wire, I) lock, J) multiple-protections.

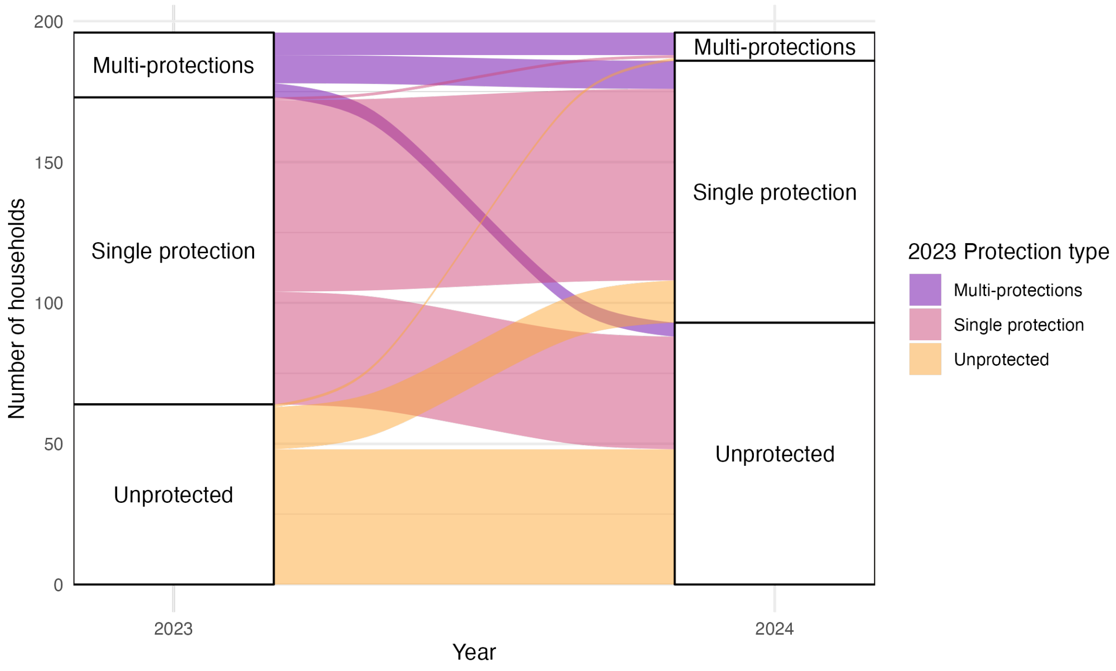


Figure S2: Year over year changes in protection type (unprotected, single protection, multiple protections).

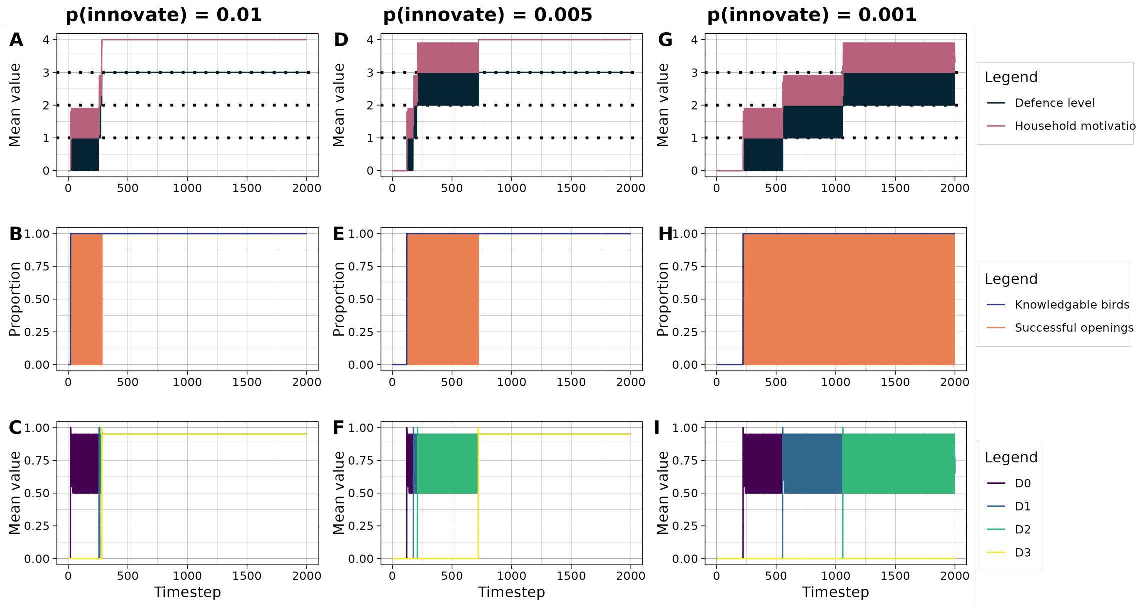


Figure S3: Effects of innovation rate (1 household vs. 1 cockatoo).

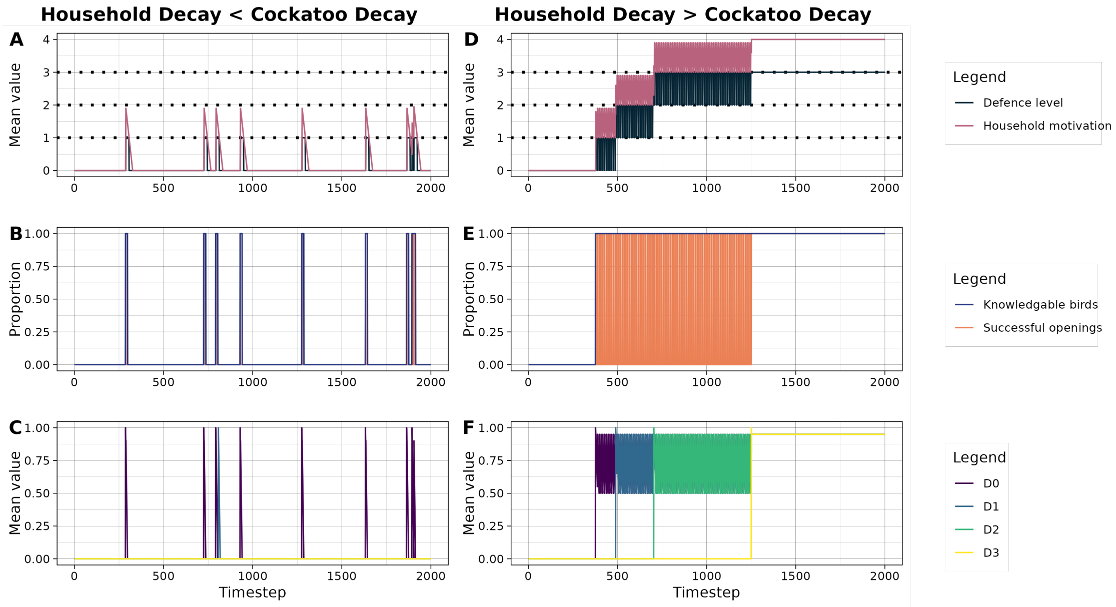


Figure S4: Effects of decay rates (1 household vs. 1 cockatoo).

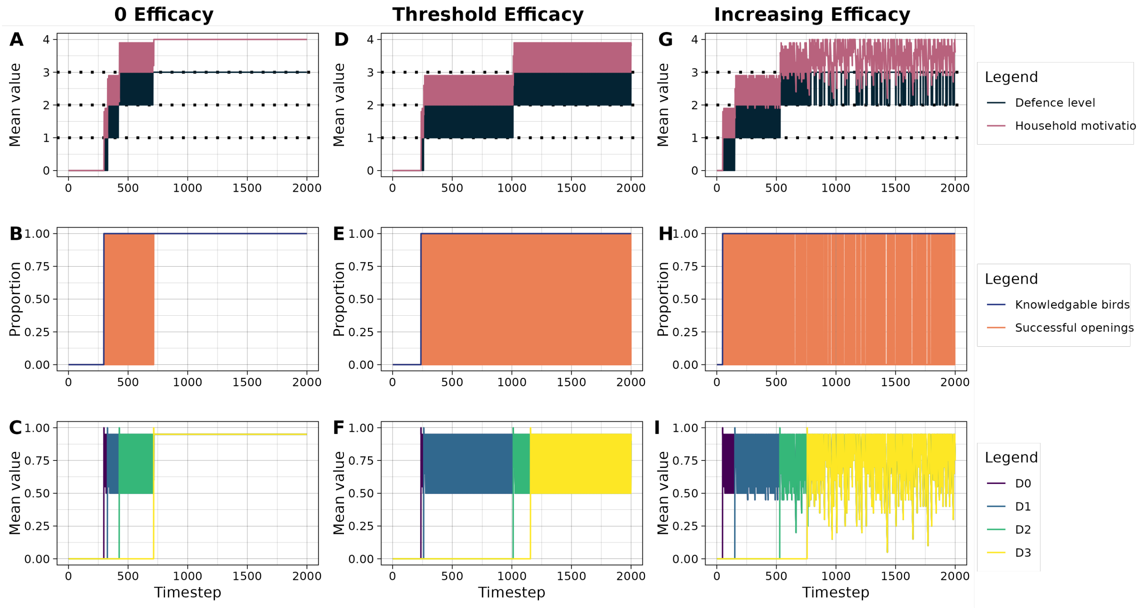


Figure S5: Effects of defence efficacy (1 household vs. 1 cockatoo).

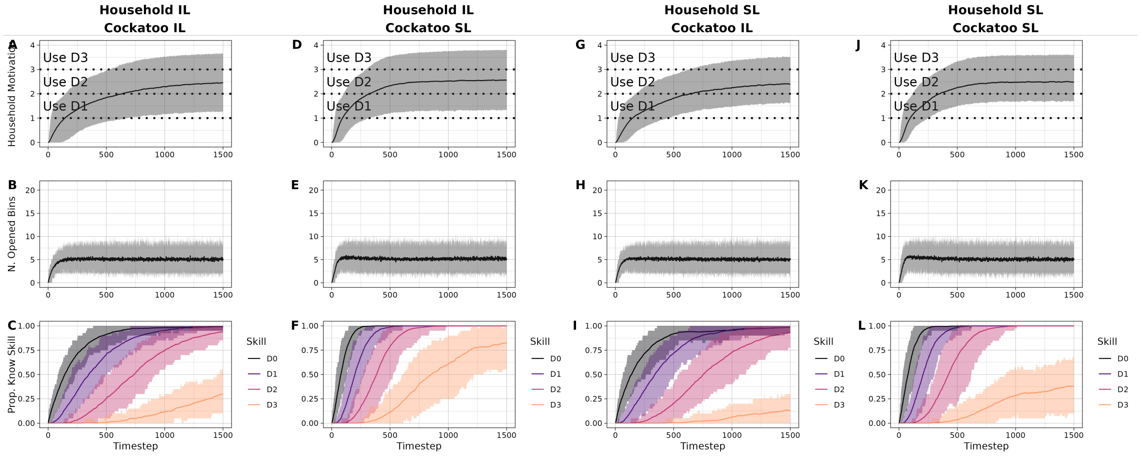


Figure S6: Simulation dynamics of different learning types under random movement.

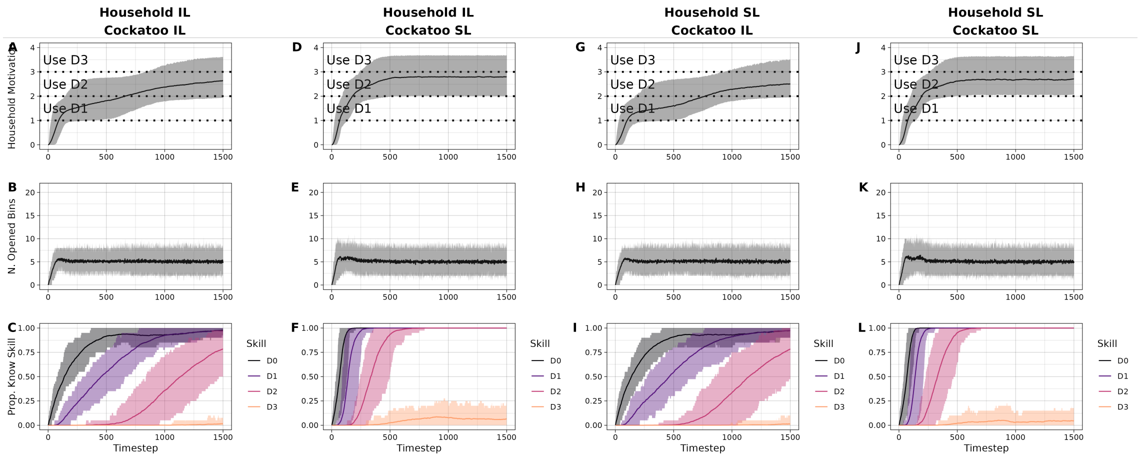


Figure S7: Simulation dynamics of different learning types under defence sensitive movement.

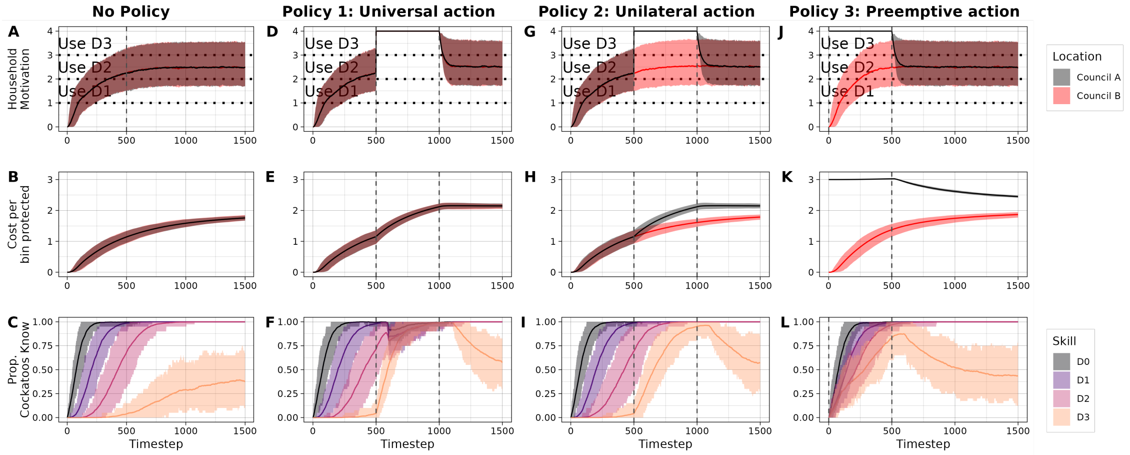


Figure S8: Simulation dynamics of different policies under random movement.

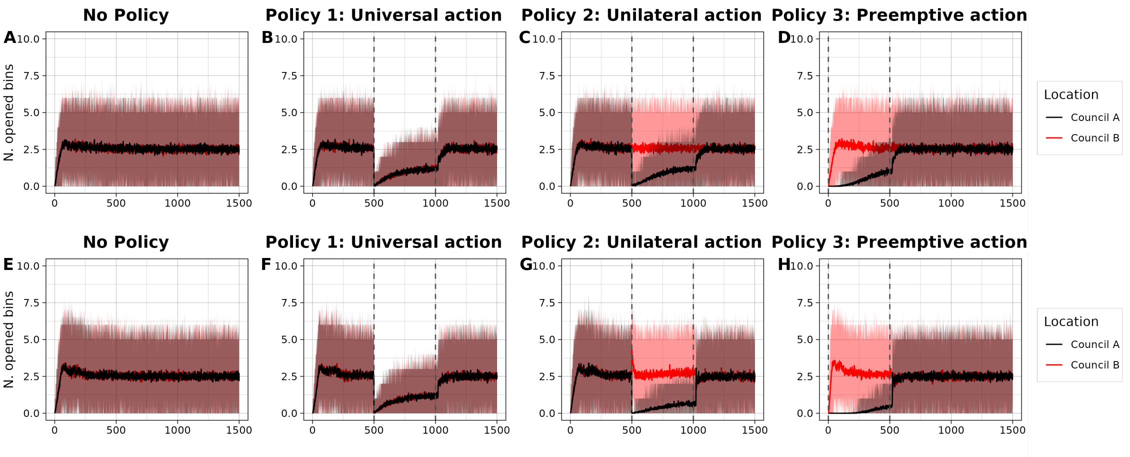


Figure S9: Number of bins open under different policies. A,B,C,D) Random movement. E,F,G,H) Defense sensitive movement.

| Method | N. bins | | Prop. Bins | | N. addresses | | Prop. Addresses | |
|---|---|---|---|---|---|---|---|---|
| | 2023 | 2024 | 2023 | 2024 | 2023 | 2024 | 2023 | 2024 |
| rim | 2 | 0 | 0.003 | 0.000 | 0 | 0 | 0 | 0.000 |
| lock | 121 | 67 | 0.165 | 0.098 | 91 | 64 | 0.236 | 0.141 |
| rope | 25 | 12 | 0.034 | 0.017 | 18 | 12 | 0.047 | 0.026 |
| hook | 6 | 5 | 0.008 | 0.007 | 2 | 4 | 0.005 | 0.009 |
| time | 3 | 0 | 0.004 | 0.000 | 1 | 0 | 0.003 | 0.000 |
| wire | 1 | 5 | 0.001 | 0.007 | 0 | 5 | 0 | 0.011 |
| stick | 36 | 47 | 0.049 | 0.068 | 31 | 42 | 0.080 | 0.093 |
| block | 5 | 3 | 0.007 | 0.004 | 4 | 3 | 0.010 | 0.007 |
| weight | 54 | 34 | 0.074 | 0.049 | 42 | 31 | 0.109 | 0.068 |
| bottle | 33 | 33 | 0.045 | 0.048 | 22 | 30 | 0.057 | 0.066 |
| spring | 4 | 18 | 0.005 | 0.026 | 2 | 14 | 0.005 | 0.031 |
| bungee | 3 | 0 | 0.004 | 0.000 | 3 | 0 | 0.008 | 0.000 |
| unprotected | 407 | 445 | 0.556 | 0.648 | 145 | 232 | 0.376 | 0.512 |
| lock;rope | 1 | 0 | 0.001 | 0.000 | 1 | 0 | 0.003 | 0.000 |
| block;lock | 1 | 0 | 0.001 | 0.000 | 1 | 0 | 0.003 | 0.000 |
| lock;stick | 1 | 1 | 0.001 | 0.001 | 1 | 1 | 0.003 | 0.002 |
| bottle;lock | 4 | 1 | 0.005 | 0.001 | 3 | 1 | 0.008 | 0.002 |
| rope;weight | 2 | 0 | 0.003 | 0.000 | 2 | 0 | 0.005 | 0.000 |
| lock;spikes | 1 | 2 | 0.001 | 0.003 | 1 | 2 | 0.003 | 0.004 |
| lock;weight | 1 | 0 | 0.001 | 0.000 | 1 | 0 | 0.003 | 0.000 |
| bottle;hook | 0 | 1 | 0.000 | 0.001 | 0 | 0 | 0.000 | 0.000 |
| stick;weight | 1 | 0 | 0.001 | 0.000 | 1 | 0 | 0.003 | 0.000 |
| block;weight | 0 | 2 | 0.000 | 0.003 | 0 | 1 | 0.000 | 0.002 |
| bottle;bottle | 10 | 2 | 0.014 | 0.003 | 6 | 2 | 0.016 | 0.004 |
| bottle;weight | 1 | 0 | 0.001 | 0.000 | 0 | 0 | 0 | 0.000 |
| weight;weight | 1 | 3 | 0.001 | 0.004 | 1 | 3 | 0.003 | 0.007 |
| bottle;spikes | 0 | 1 | 0.000 | 0.001 | 0 | 1 | 0.000 | 0.002 |
| bottle;hook;lock | 1 | 0 | 0.001 | 0.000 | 1 | 0 | 0.003 | 0.000 |
| bottle;bottle;lock | 2 | 2 | 0.003 | 0.003 | 1 | 2 | 0.003 | 0.004 |
| bottle;bottle;weight | 3 | 0 | 0.004 | 0.000 | 3 | 0 | 0.008 | 0.000 |
| bottle;bottle;bottle | 0 | 1 | 0.000 | 0.001 | 0 | 1 | 0.000 | 0.002 |
| bottle;bottle;hook;lock | 0 | 1 | 0.000 | 0.001 | 0 | 1 | 0.000 | 0.002 |
| bottle;bottle;weight;weight | 1 | 0 | 0.001 | 0.000 | 1 | 0 | 0.003 | 0.000 |
| bottle;bottle;bottle;bottle | 0 | 1 | 0.000 | 0.001 | 0 | 1 | 0.000 | 0.002 |
| bottle;bottle;rope;weight;weight | 1 | 0 | 0.001 | 0.000 | 1 | 0 | 0.003 | 0.000 |

Table S1: Proportion of addresses using each protection method.

| Year | Protection type | N. bins | Prop. Bins | N. addresses | Prop. Addresses |
|---|---|---|---|---|---|
| 2023 | Unprotected | 407 | 0.556 | 145 | 0.376 |
| 2023 | Single protection | 293 | 0.400 | 213 | 0.552 |
| 2023 | Multi-protections | 32 | 0.044 | 28 | 0.073 |
| 2024 | Unprotected | 445 | 0.648 | 232 | 0.512 |
| 2024 | Single protection | 224 | 0.326 | 205 | 0.453 |
| 2024 | Multi-protections | 18 | 0.026 | 16 | 0.035 |

Table S2: Summary of each protection type by year.

| Year | Measure | Observed | P-Value |
|---|---|---|---|
| **driving distance** | | | |
| 2023 | is protected | 0.1740318 | 0.000 |
| 2024 | is protected | 0.0255443 | 0.209 |
| 2023 | efficacy | 0.1673295 | 0.000 |
| 2024 | efficacy | 0.0053983 | 0.381 |
| 2023 | protection method | 0.1116068 | 0.000 |
| 2024 | protection method | 0.0556224 | 0.001 |
| **geodesic distance** | | | |
| 2023 | is protected | 0.1132614 | 0.001 |
| 2024 | is protected | 0.0208968 | 0.184 |
| 2023 | efficacy | 0.1222073 | 0.000 |
| 2024 | efficacy | 0.0156535 | 0.250 |
| 2023 | protection method | 0.0778326 | 0.000 |
| 2024 | protection method | 0.0327297 | 0.006 |

Table S3: Bin protection assortativity values for 2023 and 2024. Columns include the type of measurement, the year, the observed assortativity, and a p-value obtained from 1000 permutation tests. Assortativity was measured both on inverse geodesic distance and inverse driving distance networks.

| ID | Age | Sex | Unprotected | Block | Loose weight | Bottle | Stick | Rope | Spring | Fixed weight | Lock |
|---|---|---|---|---|---|---|---|---|---|---|---|
| RLO | A | M | 5/5 | - | 1/1 | 0/1 | - | - | 0/1 | - | 0/1 |
| ROL | U | U | 3/4 | - | - | - | - | - | - | 0/1 | 0/2 |
| VNV | A | M | 2/4 | - | - | - | - | - | - | - | - |
| VPO | U | U | 2/4 | - | - | 0/1 | - | - | - | - | - |
| ORG | U | M | 2/3 | - | - | - | - | - | - | - | 1/1 |
| RLG | J | U | 2/2 | - | - | - | - | - | - | - | - |
| VPN | A | M | 1/5 | - | - | - | 0/2 | - | - | - | 0/1 |
| VRB | A | M | 1/2 | - | (1)/1 | - | - | - | - | - | - |
| OGV | U | U | 1/1 | - | - | 1/2 | 0/1 | - | 0/1 | - | 1/2 |
| VON | A | M | 1/1 | - | - | - | - | - | - | - | - |
| NVL | U | U | 0/2 | - | - | - | - | - | - | - | - |
| RNGP | U | M | 0/2 | - | - | - | - | - | - | - | 0/1 |
| ORV | U | U | 0/1 | - | - | - | - | - | - | - | - |
| RBG | A | M | 0/1 | - | - | - | - | - | - | - | - |
| RGL | A | F | 0/1 | - | - | - | - | - | - | - | - |
| RVG | U | M | 0/1 | - | - | - | - | - | - | - | - |
| VBO | A | M | 0/1 | - | - | - | 0/2 | - | - | 0/2 | 0/3 |
| VBP | A | M | 0/1 | - | - | - | - | - | - | - | - |
| VPBR | J | M | 0/1 | - | - | - | - | - | - | - | - |
| VPR | U | M | 0/1 | - | - | - | - | - | - | - | 0/1 |
| NLN | U | U | - | 1/1 | - | - | - | 0/1 | - | - | - |
| NPO | U | U | - | - | - | - | 0/2 | - | - | - | - |
| NPV | J | U | - | - | - | - | - | - | - | - | 0/1 |
| NVN | U | U | - | - | - | - | - | - | - | - | 0/1 |
| RGBR | U | F | - | - | 0/1 | - | - | - | - | - | - |
| RGR | U | U | - | - | - | - | - | 0/1 | - | - | - |
| RNP | U | F | - | - | - | - | - | - | 0/2 | - | - |
| RPG/PRG | A | M | - | - | 0/1 | - | 0/1 | - | - | - | - |
| VGL | A | F | - | 0/1 | - | - | - | - | - | - | - |
| VOL | J | F | - | - | - | - | - | 0/1 | - | - | - |
| VPL | A | M | - | - | 0/1 | - | 0/1 | - | - | - | - |

Table S4: Individual-level summary of bin-opening transects. Reported are the ID, age, sex and number of successfully opened / number of attempts. We not that bird VRB was observed defeating, but not fully opening a loose weight defended bin.

| Defence type | N birds | N birds attempted | N Attempts | N birds defeated defence | N birds opened bin |
|---|---|---|---|---|---|
| unprotected | 109 | 29 | 117 | 9 | 9 |
| weight | 104 | 13 | 54 | 13 | 6 |
| bottle | 89 | 16 | 73 | 2 | 1 |
| lock | 100 | 20 | 57 | 2 | 0 |
| Total Unique | 153 | 48 | 301 | 14 | 11 |

Table S5: Summary of knowledge assay. Columns are the defence type, arranged by difficulty, the total number of attempts, the number of unique birds to attempt, the number of birds that successfully defeated the defence, and the number of birds who then went on to open the bin.

| Bird ID | Unprotected | Weight | Bottle | Lock |
|---|---|---|---|---|
| RLG | 19 (19) / 19 | 11 (11) / 11 | 0 (1) / 8 | 0 (0) / 12 |
| VPR | 10 (10) / 10 | 6 (7) / 7 | 0 (0) / 1 | - |
| ROL | 6 (6) / 6 | - | - | - |
| NRO | 4 (4) / 6 | - | - | - |
| VPN | 3 (3) / 5 | - | - | - |
| NGO | 2 (2) / 2 | 2 (2) / 2 | - | - |
| VLG | 2 (2) / 2 | - | - | - |
| NGL | 1 (1) / 1 | - | - | - |
| RLO | 1 (1) / 1 | 7 (7) / 7 | - | 0 (0) / 2 |
| N | 0 (0) / 1 | - | - | - |
| NPN | 0 (0) / 2 | - | - | - |
| NRV | 0 (0) / 1 | 0 (4) / 4 | - | 0 (0) / 2 |
| OGV | - | - | - | 0 (0) / 2 |
| OLN | - | - | 0 (0) / 1 | - |
| ORB | 0 (0) / 1 | - | - | - |
| ORG | - | 0 (1) / 1 | - | - |
| ORP | 0 (0) / 2 | - | - | - |
| ORV | 0 (0) / 1 | - | - | - |
| RBG | - | 0 (1) / 1 | - | - |
| RBP | - | - | - | 0 (0) / 1 |
| RGB | - | - | 0 (0) / 1 | - |
| RGBR | 0 (0) / 1 | - | - | - |
| RLP | - | - | 0 (0) / 1 | - |
| RNGP | 0 (0) / 6 | - | 0 (0) / 7 | 0 (0) / 2 |
| RNV | - | 0 (1) / 1 | - | - |
| RON | - | - | - | 0 (0) / 1 |
| RPG/PRG | 0 (0) / 1 | - | - | - |
| RPO | - | - | - | 0 (0) / 1 |
| RVBL | 0 (0) / 4 | - | - | - |
| RVN | - | - | 0 (0) / 1 | - |
| VBL | 0 (0) / 1 | 3 (3) / 3 | 1 (2) / 16 | 0 (0) / 8 |
| VBO | - | - | 0 (0) / 8 | 0 (0) / 3 |
| VBP | 0 (0) / 10 | - | - | - |
| VGB | - | - | 0 (0) / 3 | 0 (1) / 6 |
| VGL | - | - | 0 (0) / 1 | - |
| VGV | 0 (0) / 3 | - | - | - |
| VLV | - | - | 0 (0) / 1 | 0 (0) / 1 |
| VNP | - | - | - | 0 (0) / 1 |
| VNV | 0 (0) / 7 | 0 (4) / 4 | - | 0 (0) / 4 |
| VOB | 0 (0) / 1 | - | - | - |
| VON | 0 (0) / 2 | 9 (9) / 9 | 0 (0) / 14 | 0 (0) / 1 |
| VPBR | - | - | 0 (0) / 2 | - |
| VPL | 0 (0) / 7 | 0 (1) / 1 | - | 0 (1) / 2 |
| VPO | 0 (0) / 1 | - | - | - |
| VRB | 0 (0) / 5 | 0 (3) / 3 | - | 0 (0) / 2 |
| VRL | - | - | 0 (0) / 4 | 0 (0) / 1 |
| VRNR | - | - | - | 0 (0) / 1 |
| VRP | 0 (0) / 8 | - | 0 (0) / 4 | 0 (0) / 4 |

Table S6: **Individual-level summary of knowledge assay.** Columns are bird identity, and for each defence level, the values are N Completed (N Defeated) / N Attempted. Completed: the number of times the bin was opened after the defence was defeated, Defeated the number of times the protection was defeated, and Attempted: the number of times the task was attempted regardless of success.

| ID | Age | Sex | Total skills | Open | Block | Weight | Bottle | Lock |
|---|---|---|---|---|---|---|---|---|
| OGV | U | U | 3 | TRUE | FALSE | FALSE | TRUE | TRUE |
| ORG | U | M | 3 | TRUE | FALSE | TRUE | FALSE | TRUE |
| RLG | J | U | 3 | TRUE | FALSE | TRUE | TRUE | FALSE |
| VBL | A | M | 3 | TRUE | FALSE | TRUE | TRUE | FALSE |
| NGO | A | M | 2 | TRUE | FALSE | TRUE | FALSE | FALSE |
| RLO | A | M | 2 | TRUE | FALSE | TRUE | FALSE | FALSE |
| VNV | A | M | 2 | TRUE | FALSE | TRUE | FALSE | FALSE |
| VON | A | M | 2 | TRUE | FALSE | TRUE | FALSE | FALSE |
| VPR | U | M | 2 | TRUE | FALSE | TRUE | FALSE | FALSE |
| VRB | A | M | 2 | TRUE | FALSE | TRUE | FALSE | FALSE |
| NLN | U | U | 2 | TRUE | TRUE | FALSE | FALSE | FALSE |
| NGL | J | U | 1 | TRUE | FALSE | FALSE | FALSE | FALSE |
| NRO | U | U | 1 | TRUE | FALSE | FALSE | FALSE | FALSE |
| ROL | U | U | 1 | TRUE | FALSE | FALSE | FALSE | FALSE |
| VLG | A | F | 1 | TRUE | FALSE | FALSE | FALSE | FALSE |
| VPN | A | M | 1 | TRUE | FALSE | FALSE | FALSE | FALSE |
| VPO | U | U | 1 | TRUE | FALSE | FALSE | FALSE | FALSE |
| VPL | A | M | 2 | FALSE | FALSE | TRUE | FALSE | TRUE |
| NRV | J | U | 1 | FALSE | FALSE | TRUE | FALSE | FALSE |
| RBG | A | M | 1 | FALSE | FALSE | TRUE | FALSE | FALSE |
| RNV | U | U | 1 | FALSE | FALSE | TRUE | FALSE | FALSE |
| VGB | A | M | 1 | FALSE | FALSE | FALSE | FALSE | TRUE |

Table S7: **Individual-level summary of assay and transect data combined.** These are the 22 SCC we observed as having some knowledge about the bins. Columns include ID, age, sex, whether they were observed opening, and which protections they defeated.
